# The Goldilocks Window of Personalized Chemotherapy: An Immune Perspective

**DOI:** 10.1101/495184

**Authors:** Derek S. Park, Mark Robertson-Tessi, Philip K. Maini, Michael B. Bonsall, Robert A. Gatenby, Alexander R. A. Anderson

**Author notes:** Corresponding Authors: Derek Park, 12902 USF Magnolia Drive, Mailstop: SRB 4, Tampa, FL 33612, Alexander R. A. Anderson, 12902 USF Magnolia Drive, Mailstop: SRB 4, Tampa, FL 33612.

## Abstract

The immune system is increasingly being recognized for its untapped potential in being recruited to attack tumors in cancer therapy. The main challenge, however, is that most tumors exist in a state of immune tolerance where the patient’s immune system has become insensitive to the cancer cells. In order to investigate the ability to use chemotherapy to break immune tolerance, we created a mathematical modeling framework for tumor-immune dynamics. Our results suggest that optimal chemotherapy scheduling must balance two opposing objectives: maximal tumor reduction and preserving patient immune function. Successful treatment requires therapy to operate in a ‘Goldilocks Window’ where patient immune health is not overly compromised. By keeping therapy ‘just right’, we show that the synergistic effects of immune activation and chemotherapy can maximize tumor reduction and control.

**Statement of Significance:** In order to maximize the synergy between chemotherapy and anti-tumor immune response, lymphodepleting therapy must be balanced in a ‘Goldilocks Window’ of optimal dosing.

## Introduction

By the time a tumor is clinically detectable, it is no longer subject to significant anti-tumor response from the innate and acquired components of the host immune system. Mechanistically, this immune tolerance is the result of complex interactions among tumor cells, T cells, and secreted cytokines [1]. CD8+ effector T cells, also known as cytotoxic T lymphocytes (CTLs), are an important component of the adaptive immune system that responds to tumor antigens and induces cell death.

A major barrier to effective CTL response in tumors is suppression by T regulatory cells (Tregs), which inhibit CTL cytotoxic activity via cell-cell contact ([2], [3]) as well as through secreted factors such as TGF-beta ([4] [5]). They have posed challenges for cancer immunotherapies as well as preventing the activation of the immune system during more traditional therapy approaches ([3], [6]). Tregs also appear to play a critical role in limiting immune response in maternal tolerance of the fetus and protection of commensal bacteria from the host immune system [2].

Multiple methods have been investigated to break the immune system from tolerance and revive anti-tumor immune activity. The initial focus of these approaches included activation of CTLs through immunostimulatory cytokines such as interleukin-2 (IL-2). More recently, lymphodepleting chemotherapy has been recognized to have paradoxical but important immunostimulatory effects. Heavy lymphodepletion has been reported to enhance the impact of adoptively transferred tumor-specific T cells ([7]). This leads to the interesting question of whether or not lymphodepletion can also enhance the efficacy of existing T-cell populations to mount an anti-tumor response. While Gemcitabine, 5-Fluorouracil and other cytotoxic drugs can initially suppress immune subpopulations, notably B and T cells, the subsequent proliferation of the immune cells when therapy is completed provides a transient period in which immune response to tumor antigens can be restored. An obvious question then arises: is there a better chemotherapy schedule that could maximize tumor kill and also enhance immune response?

To investigate the dynamics of this transient immune response following chemotherapy, we created a mathematical model of the complex tumor-immune dynamics that occur during multiple cycles of chemotherapy. In particular, we investigated three, clinically-relevant, therapeutic dynamics: immunodepletion, immunostimulation via vaccination, and immunosupportive prophylactics. We identified significant immune trade-offs during chemotherapy as well as the relevant patient metrics that determine the magnitude and severity of these compromises. Further, by exploring the impact of clinically-established, as well as more experimental treatment, decisions we illustrate a more complex interplay between chemotherapy and patient immune dynamics than has been previously investigated. Our results indicate that optimal chemotherapy requires identification of a ‘Goldilocks Window’ in which treatment can both induce cytotoxic effects in the tumor and enhance the immune response to tumor antigens. Therefore, instead of the one-size-fits-all paradigm of fixed therapy regimens, patient immune biology should be a key consideration when developing personalized chemotherapy strategies.

## Methods

Quick guide to equations and assumptions:

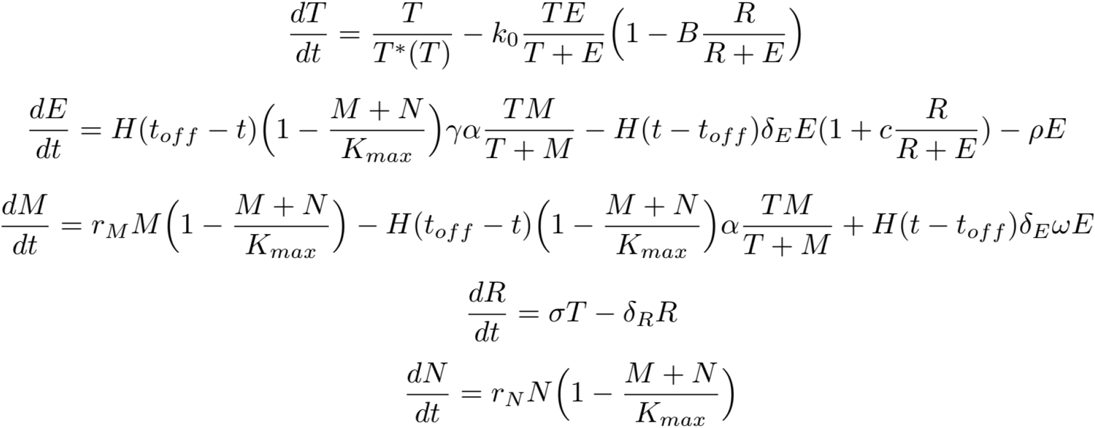

Our model assumes that tumor cells (T) grow unless checked by T effector cells (E). However, effector cells are themselves inhibited by T regulatory cells (R) that are recruited at a rate σ by tumor antigens. This leads to effector-cell-mediated tumor cell death being moderated by the quantity of T regulatory cells 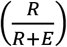. Effector cells exhibit different behaviors during immune expansion and immune contraction. This switching behavior is modeled with the Heaviside function (*H*(*t_off_* − *t*)). During the immune expansion phase, effector cells are recruited based on both available memory cells (M) and the tumor burden 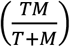. Memory cells are the pool of T cells from which effector cells are derived. During immune expansion, the antigenicity of the tumor (*α*) induces differentiation to effector cells 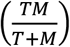. However, as immune tolerance sets in, there is a contraction in the effector T cell population. This is caused by degradation of effector cells by T regulatory cells 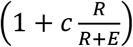. During immune contraction, there is also a small influx into the memory T cell compartment due to conversion of effector cells to memory T cells (*ωE*). Finally, the total lymphocyte population is represented by naïve cells (N) which replicate in a logistic growth model 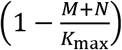.

### Overall Model Design

A central assumption of this work is that a clinically-detectable tumor has induced a tolerant state in which the immune system can no longer respond to tumor antigens. Chemotherapy temporarily removes this tolerance through lymphodepletion, which eliminates Tregs and allows a burst of immune response. However, the lymphodepletion itself also kills CTLs and therefore reduces the potential cytotoxic efficacy. This double-edged response to chemotherapy implies that there is an optimal therapeutic strategy. If the dose is too high, then the few remaining immune cells will not be able to take advantage of the tolerance breaking; if the dose is too low, then the immune depletion will be insufficient to break tolerance. In addition to these immune effects, the chemotherapy itself can induce cancer cell death affecting both the tumor size directly and releasing tumor antigens, adding another layer of complexity to the tumor-immune dynamics.

We develop a mathematical model that includes five major populations of cells: Tumor cells (T), T effector cells (also known as cytotoxic T lymphocytes, CTLs, and denoted as E), T regulatory cells (Tregs, R), Memory T cells (M), and Naive T cells (N). Immune function is separated into two distinct temporal stages relative to the time of application of each chemotherapy cycle: 1) a period of CTL expansion in a sensitized immune system, immediately following the application of chemotherapy (Figure 1, panel A), and 2) CTL contraction as tolerance returns (Figure 1, panel B). The transition between these expansion and contraction phases is governed by mechanisms that remain poorly characterized, but empirically occurs 5-10 days after the expansion starts [8]. In the model, the transition time is set to 5 days after the start of the immune expansion phase. Therefore, there is a window of 5 days immediately following each cycle of chemotherapy in which the immune system is sensitive, and outside of these periods, it is tolerant.

**Figure 1:**
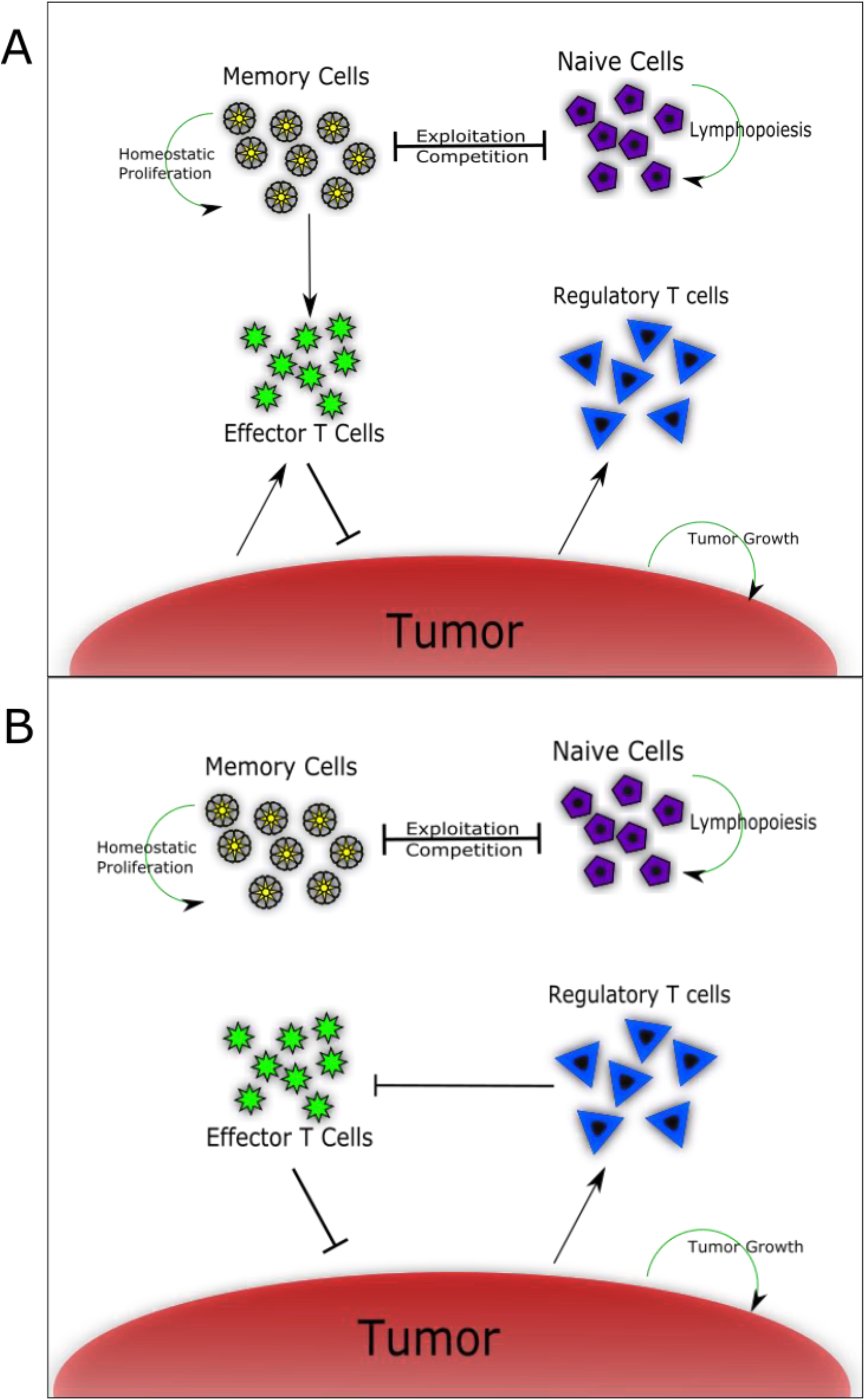
Tumor-immune dynamics during the sensitive (A) and tolerant (B) stages of the immune response. During antigen-sensitive immune expansion, CTLs are recruited from memory cells to attack tumor cells. Tregs are being recruited but have not yet started significantly inhibiting CTL responses. During immune contraction once tolerance sets in, Tregs exert an active inhibitory pressure on CTLs. Expansion of memory cells into CTLs ceases. Both stages of the immune response are characterized by competition between memory and naïve immune cells for common cytokine pools as well as homeostatic proliferation and lymphopoiesis.

During the phase in which the immune system is sensitive to the tumor, a few key processes occur. CTLs, which target and kill the tumor, are recruited from a memory cell population due to detection and response to tumor antigens [8]. These memory cells are constantly undergoing a low level of replenishing proliferation, but they only convert to CTLs during the sensitive expansion phase following lymphodepletion. During this phase, there is also tumor-mediated recruitment of Tregs. This eventually causes a significant shift in immune dynamics, leading to a contraction of the effector compartment during the tolerized phase. Under tolerance, there is no longer a significant recruitment of effector cells from the memory cell compartment. Instead, while the existing effector cells do carry out some tumor-killing function, the Tregs decrease the CTL number.

### Tumor dynamics

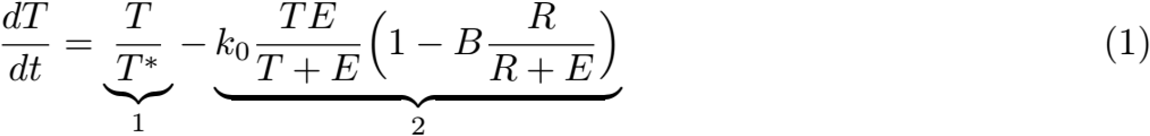

Tumor growth dynamics are approximated via a combination of exponential growth for smaller tumors and power law growth for larger tumors, as shown in the first term on the right hand side of Eq. (1). The transition between these growth dynamics is governed largely by the *T*\* term as defined in equation (2), following the implementation of tumor-immune growth dynamics described in [9],

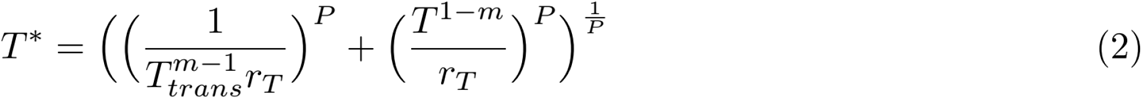

*T*\* employs the method of modeling tumor growth in [9] (specifically the first term on the right hand side of equation 1) by having tumor populations transition from exponential to power law growth. As the authors note, tumors are not able to sustain early exponential growth due to physical and nutrient limitations. A more appropriate model is where there is exponential growth early which then transitions to a power law growth at larger tumor sizes. The size at which this transition in growth occurs is *T_trans_* and the smoothness of this transition is governed by the exponent P. The growth term *r_T_* represents the growth rate and how aggressively the tumor is developing.

The second term of Eq. (1) on the right hand side represents the tumor loss due to killing by CTLs. The parameter *k*_0_ represents the CTL cytotoxic efficacy, with the actual tumor kill rate being dependent upon the relative numbers of tumor and effector cells 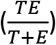. However, this rate is mitigated by the presence of Tregs, with *B* representing their inhibition efficacy. As Tregs increase in density, the CTL-mediated tumor death rate decreases.

### Effector T cell dynamics

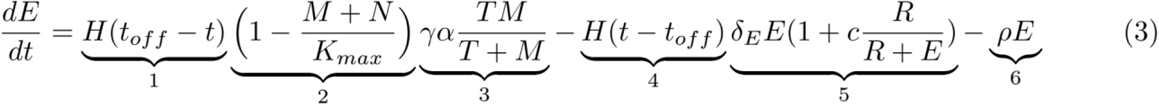

CTL dynamics are modeled in two phases, expansion (terms 1-3) and contraction (terms 4-6), as described above. Terms 1 and 4 switch between these phases via the Heaviside function, with time *t_off_* being the length of the expansion phase (5 days) immediately following each round of chemotherapy. Terms 2 and 3 chiefly govern the growth of CTLs during immune sensitivity to the tumor. CTLs are generated based upon the antigenicity of the tumor (*α*) as well as the number of tumor and memory T cells. The antigenicity describes how much of an immune response is promoted by the tumor. Modulating this is an amplification rate, *γ*, since one memory cell can yield multiple effector cells. Term 2 represents a moderating term where there is a maximum number of memory and naïve lymphocytes that can be supported by the cytokine pool. This general paradigm of effector cell function being limited by cytokine availability has been supported by lymphodepletion studies that have shown increased CTL activity when IL-7 and IL-15 cytokine-responsive cells were removed. With fewer cytokine sinks, CTL activity was increased [10]. When the immune compartment is full and in homeostasis, this term will be near zero, effectively shutting down CTL recruitment; however, immediately after a dose of chemotherapy, memory and naïve T cells are depleted, which promotes CTL expansion. Term 5 represents the contraction of the effector cell compartment that occurs due to immune tolerance. There is a death rate of CTLs, *δ_E_*, which is increased by the relative fraction of Tregs that are present, 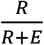. Tregs have been shown to inhibit CTLs through a variety of mechanisms, including both depriving cytokines necessary for CTL sustenance as well as direct cytolysis of CTLs [11]. Parameter c represents the suppression efficacy of Tregs. Lastly, term 6 represents the rate of conversion of effector cells back into memory cells, which is an active mechanism during immune contraction [12].

### Memory T cell dynamics

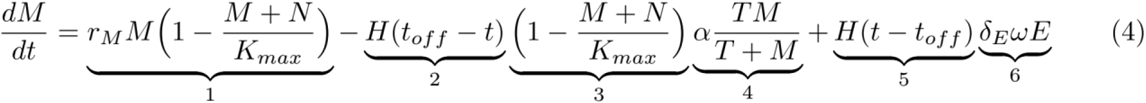

Memory cells continually replenish themselves through homeostatic growth in term 1. Parameter *r_M_* is the maximum memory T-cell growth rate, which is decreased as the memory and naïve cell numbers reach their carrying capacity, *K_max_*. During the immune expansion phase (terms 2-4), there is memory cell loss as they are converted to CTLs. The conversion rate is governed in term 4 by the relative abundances of tumor and memory cells, 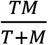, as well as the antigenicity, *α*, as mentioned above. As described in Eq. (3), the rate of recruitment is moderated by the relative homeostasis level of the overall immune system. During the contraction phase, memory cells are replenished from the CTL compartment. A fraction (*ω*) of the CTL are successfully converted back to memory cells [12]. Due to some loss and inefficiency of conversion, the fraction, *ω*, is less than the loss from the effector cell compartment, *ρ* > 0 [13].

### Regulatory T cell and naïve T cell dynamics

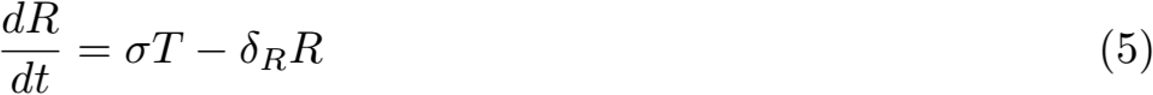

Tregs are recruited due to secretion of factors such as TGF-beta from peripheral precursor cells by tumor cells with recruitment rate *σ*, and decay with a rate *δ_R_* [14].

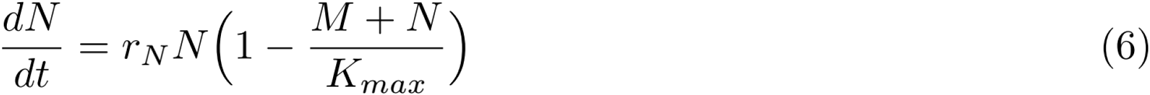

Naive T cell dynamics are largely the result of homeostatic proliferation up to a common carrying capacity of *K_max_*, which is the maximum number of memory and naïve T cells in the immune system [15]. The naive cell replenishment rate is determined by *r_N_*.

The model was parameterized based on literature sources when possible, as shown in Table 1. For many cases there was evidence of variation in parameters, as well as no clear study of each individual parameter in our model. This is, in part, due to approach to simplify, mathematically, certain processes in favor of focusing on the tumor-immune dynamics. Where possible, we have tried to make a biologically reasonable order-of-magnitude approximation. In order to address this parameter uncertainty we explicitly consider the impact of parameter variation on model results.

**Table 1:** Model parameters were estimated based upon both pre-existing models, chiefly Althaus *et al*., 2007 and Robertson-Tessi *et al*., 2012, as well as experimental studies. For most of the parameters, the literature often indicated significant variation and so order-of-magnitude approximations were made. Similarly, certain parameters were not succinctly captured in literature studies and were therefore estimated (*). We have addressed the impact of potential parameter variation through sensitivity studies (see Results).

| Parameter | Symbol | Value | Literature reference |
| --- | --- | --- | --- |
| Tumor Growth Coefficient | $r_T$ | 1000 cells<sup>-1</sup> day<sup>-1</sup> | Robertson-Tessi *et al.*, 2012 |
| Effector cell kill rate | $k_0$ | 1 day<sup>-1</sup> | Diefenback *et al.*, 2001 |
| Regulatory cell suppression efficacy | B | 0.75 | Robertson-Tessi *et al.*, 2012 |
| Tumor growth transition size | $T_{trans}$ | 10<sup>6</sup> cells | Robertson-Tessi *et al.*, 2012 |
| Power-Law growth exponent | m | 0.5 | Robertson-Tessi *et al.*, 2012 |
| Exponential to power smoothing term | P | 3.0 | Robertson-Tessi *et al.*, 2012 |
| Time till immune contraction | $t_{off}$ | 5 days | Althaus *et al.*, 2007 |
| Maximum sustainable number of effector, naïve, and memory cells | $E_{max}$ | 10<sup>12</sup> cells | Bains *et al.*, 2009 |
| Tumor antigenicity | $\alpha$ | 1* | Robertson-Tessi *et al.*, 2012 |
| Effector cell death rate (expansion) | $\rho$ | 0.0-0.1* | Vignali *et al.*, 2008 |
| Effector cell death rate (contraction) | $\delta_E$ | 0.13 | Althaus *et al.*, 2009 |
| Effector cell death rate due to regulatory T cells | c | 0.01* | Robertson-Tessi *et al.*, 2012 |
| Memory cell expansion factor | $\gamma$ | 100* | Althaus *et al.*, 2007; Arstila *et al.*, 1999 |
| Tumor-mediated regulatory cell recruitment rate | $\sigma$ | 0.01 | Antony *et al.*, 2005; Robertson-Tessi *et al.*, 2012 |
| Regulatory cell death rate | $\delta_R$ | 0.1* | Robertson-Tessi *et al.*, 2012 |
| Memory cell growth rate | $r_M$ | 0.01 day<sup>-1</sup>* | Bains *et al.*, 2009 |
| Memory cell reconversion rate | $\omega$ | 0.01* | Bains *et al.*, 2009 |
| Naïve cell growth rate | $r_N$ | $0.1 \text{ day}^{-1}$ | Bains *et al.*, 2009 |
| Maximum number of naïve T cells | $K_{\max}$ | $10^{12}$ cells | Lythe *et al.*, 2016 |
| Baseline chemotherapy strength | $C_0$ | Varied in simulation | |

### Simulating chemotherapy and evaluating outcomes

To establish tolerance in the system and allow transients from initial conditions to dampen before applying therapy, the simulation was started with a tumor size of 10^7^ cells. Chemotherapy was started when the tumor reached 10^8^ cells and was simulated as periodic doses of cytotoxic therapy at 14 day intervals (a standard cycle length). In total, 10 cycles of chemotherapy were applied. At the time of each treatment cycle, all cell populations (immune and tumor) were instantaneously reduced by a fraction representing the cytotoxic effect of chemotherapy. Immune cells were reduced by the same baseline fraction (**C**_0_) on each cycle. To account for tumor resistance to therapy, the fractional tumor reduction for cycle **i** (**C_i_**) was linearly reduced with each cycle, such that the cytotoxic fraction on the last cycle was 75% of **C**_0_. Approximating the impact of chemoresistance on drug efficacy is challenging since values vary for different classes of drugs. To further complicate resistance impacts, Hao et al. in [16] noted dose-dependent differences between resistant and resensitized prostate cancer cell populations to docetaxel (Figure 2, Panel A). The relative advantage of resistant to sensitive cells varied from almost nothing (at very low doses) to a 400% difference. The value of 75% chemotherapy efficacy at resistance represents a 33% advantage of survivorship for a resistant population versus a susceptible population. It is a conservative estimate of the impact of resistance, but we believe it is reasonable given that tumor populations are unlikely to be entirely homogeneously resistant. Varying this range is a relevant question for future research. For our purposes, **C_i_** is given by:

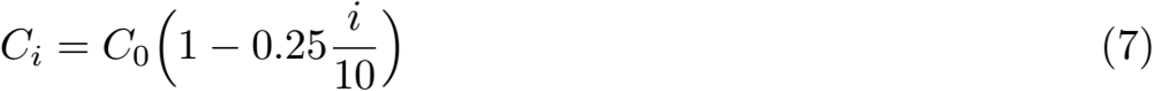

**Figure 2:**
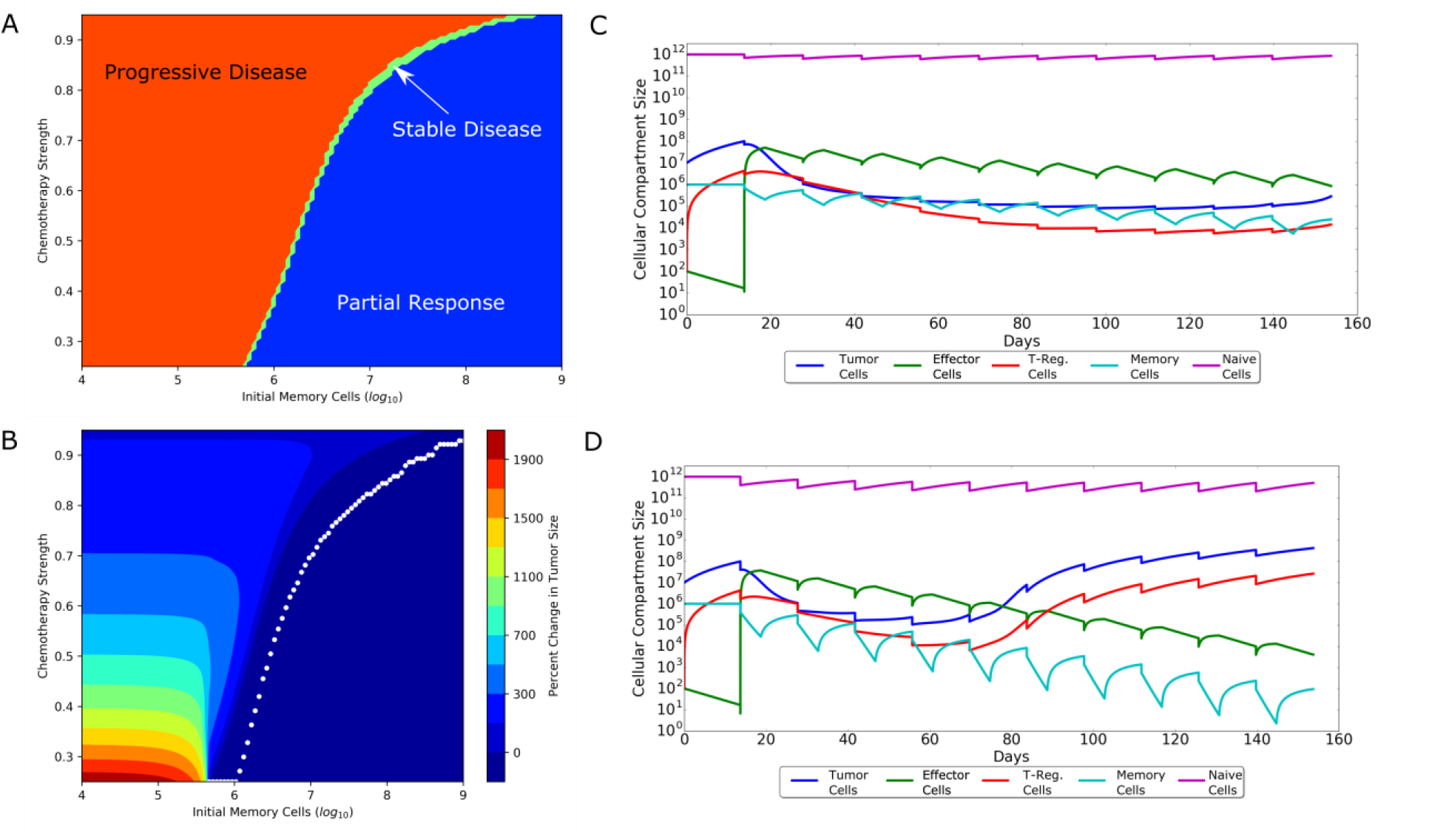
Interaction of memory-cell populations and chemotherapy strength on treatment outcomes. RECIST outcomes are shown in panel A with progressive disease (red), stable disease (yellow), partial response (light blue) and complete response (dark blue). (B) Finer grade responses are shown as percent changes in tumor size after therapy versus the initial starting size (10^8^ cells). The underlying dynamic reasons for these differences can be seen in the memory populations during low (C) and high dose chemotherapy (D). Low dose chemotherapy allows memory populations (light blue) to be sustained for longer and generate larger CTL responses (green). High dose chemotherapy, however, depletes memory cells faster and leads to declining CTL responses and concurrent tumor escape.

The final tumor size after 10 cycles of chemotherapy was compared to the tumor size at the start of treatment (10^8^ cells) and evaluated according to RECIST categories. Specifically, a total loss of tumor (<−99% change in size) is a complete response (CR). A change between −30% and −99% is considered a partial response (PR). Tumor changes between −30% and +20% are classified as stable disease (SD) and changes of greater than +20% are seen as progressive disease (PD) [17]. While there are many different methods of measuring therapy efficacy impact on disease, RECIST categories were chosen here since they have correlated well with overall survival in patients across a variety of cancers.

### Simulation environment

The model was programmed in the Python language (ver. 2.7.11). The open-source packages Scipy (ver. 0.17.0), Numpy (ver. 1.10.4), and Matplotlib (ver. 1.5.1) were used for simulation of the ODEs as well as visualization of the results. The platform for the program was both an Intel(R) Core (TM) i7-6820 HQ processor as well as the high performance computing cluster at Moffitt Cancer Center, Tampa, Florida, USA.

## Results

### Influence of patient memory cell populations

To analyze the effect of the memory T-cell population on therapy, varying doses of chemotherapy were simulated for a range of memory cell population sizes. The size of the memory T-cell population at the time of therapy was a significant factor affecting the optimal therapeutic response. Memory cell population sizes are variable among patients; Arstila et al. (1999) have estimated there to be 10^6^ – 10^7^ memory T cell clones in the human body with approximately 10^5^ memory T cells per antigen [9, 18]. However, due to antigen responses being polyclonal, this suggests multiple orders of magnitude of potential variation in memory T-cell numbers. Patient memory-cell numbers influence the maximum chemotherapy dose strength before treatment failure (Figure 2). Generally, there is a minimum memory-cell population size that is necessary for any given strength of chemotherapy to be successful. Above this threshold, the more memory cells there are, the better the improvement with stronger doses of therapy. Conversely, this means that when memory-cell populations are close to the minimum threshold, chemotherapy should be similarly weak if a more favorable treatment outcome is desired. If memory cells are below the minimum threshold, then the optimal strategy is to use strong chemotherapy (Figure 2, panel A and B). This treatment solely relies on chemotherapeutic cytotoxicity with no immune stimulation.

The double-edged nature of chemotherapy on the immune system can be better understood through the transient dynamics during therapy (Figure 2, panel C and D). In cases with stronger chemotherapy dosing, there is an early decrease in tumor population levels as the cytotoxic strength of the therapy comes to bear on cancer populations. However, we observe a trend in that these therapies tend to lead to failure and larger final tumor sizes than if treated with a ‘weaker’ chemotherapy regimen. Weaker chemotherapy regimens exert lower cytotoxic burdens on the tumor but maintain tumor size reduction for the duration of therapy.

This counterintuitive result stems from the fact that cytotoxicity alone is insufficient for suppressing tumor growth, especially due to the accumulating chemoresistance. Rather, it is the synergistic effect of cytotoxicity as well as the breaking of immune tolerance and consequent recruitment of CTLs that keeps tumor populations in check. Our *in silico* treatments consistently show that there is an inherent disadvantage to high-dose chemotherapy. There is a gradual decrease in the CTL population over multiple rounds of treatment due to the net loss that stronger dosing causes in memory T-cell populations. It is these memory cells that are affected the most by chemotherapy since they can only recover relatively slowly. If the cytotoxic pressure on memory cells is greater than the recovery rate of that compartment, then even with a resensitized immune system, expansion will lead to fewer CTLs and ultimate treatment failure. In contrast, if the immunodepleting side effects of chemotherapy can be balanced with immune recovery, then more sustainable treatment responses are possible. In short, there is a tradeoff between having chemotherapy strong enough to sufficiently break tolerance, but mild enough to leave sufficient memory T cells for adequate CTL expansion. Akin to the story of Goldilocks and the three bears, the balancing of these two immunological goals leads to an intermediary chemotherapy strength that is ‘just right’. *In silico* simulation shows that this “Goldilocks Window” is highly dependent upon patient-specific, pre-existing memory T-cell populations.

### The impact of CTL efficacy

We sought to identify other relevant patient-specific immune parameters by studying the effect of CTL killing efficacy (*k*_0_). With memory-cell sizes set at 10^6^ cells, the cytotoxicity rate was varied around the biologically realistic parameter of 0.9 per day [19]. Unsurprisingly, CTL efficacy is a significant determinant of treatment success (Fig. 1). Furthermore, CTL efficacy dramatically impacts optimal chemotherapy dosing. Lower rates of CTL-mediated tumor cell death require weaker chemotherapy for more favorable treatment outcomes. As before, the underlying dynamics demonstrate the importance of a large enough memory-cell pool over the course of therapy to supply the CTL pool in sufficient numbers. With a lower value of *k*_0_, more CTLs are necessary to exert the same degree of immune control over the tumor. This in turn, necessitates a larger pool of memory T cells. Strong chemotherapy on a system with lower k_0_ values would prevent sufficient CTL expansion by rapidly diminishing the memory-cell populations. This is counterintuitive since an initial motivation may suggest that, in a situation where a patient has a weaker immune system to combat the cancer, the chemotherapy should be increased in order to compensate. However, our model suggests that the lymphodepleting impact of heavy chemotherapy on an already weaker immune system will only worsen outcomes.

### Impact of tumor growth rates

Tumor growth rates are variable, and in the model we used a value of r_T_ = 1000 cell^−1^ per day, putting growth at a doubling time of 1 day during the fastest exponential growth phase. Experimental and model analyses have shown that selection pressures on growing tumors can lead to significant heterogeneity in metabolism and growth rates [20]. Analysis of the model with different tumor growth rates revealed that optimal dosing was dependent on this variation (Fig. 3). For slower growing tumors, greater doses can be used because chemotherapeutic cytotoxicity is sufficient for controlling tumor growth. For faster growing tumors (larger r_T_) it becomes necessary to decrease chemotherapeutic strength in order to achieve optimal outcomes; chemotherapeutic cytotoxicity is insufficient alone and so CTL-mediated tumor death is necessary. Greater CTL involvement, though, imposes the same trade-off as above, in that dosing must be weakened in order to sustain memory cell populations. Importantly, for the most aggressively growing tumors, there is actually a ‘worst-case scenario’ of intermediary chemotherapy strength. Here, the worst chemotherapy is not, in fact, the strongest possible dose and is instead a ‘mid-range’ strength in treatment. At this chemotherapeutic strength, the drug alone is insufficient to cause a reduction in tumor size. However, the dose is still strong enough to lead to severe memory cell population depletion and undermines any immune efforts at constraining tumor growth. These considerations demonstrate how the tumor growth rate is a primary determinant of tumor control and, depending on the individual patient’s tumor, determines which dynamics are capable of leading to successful treatment responses.

**Figure 3:**
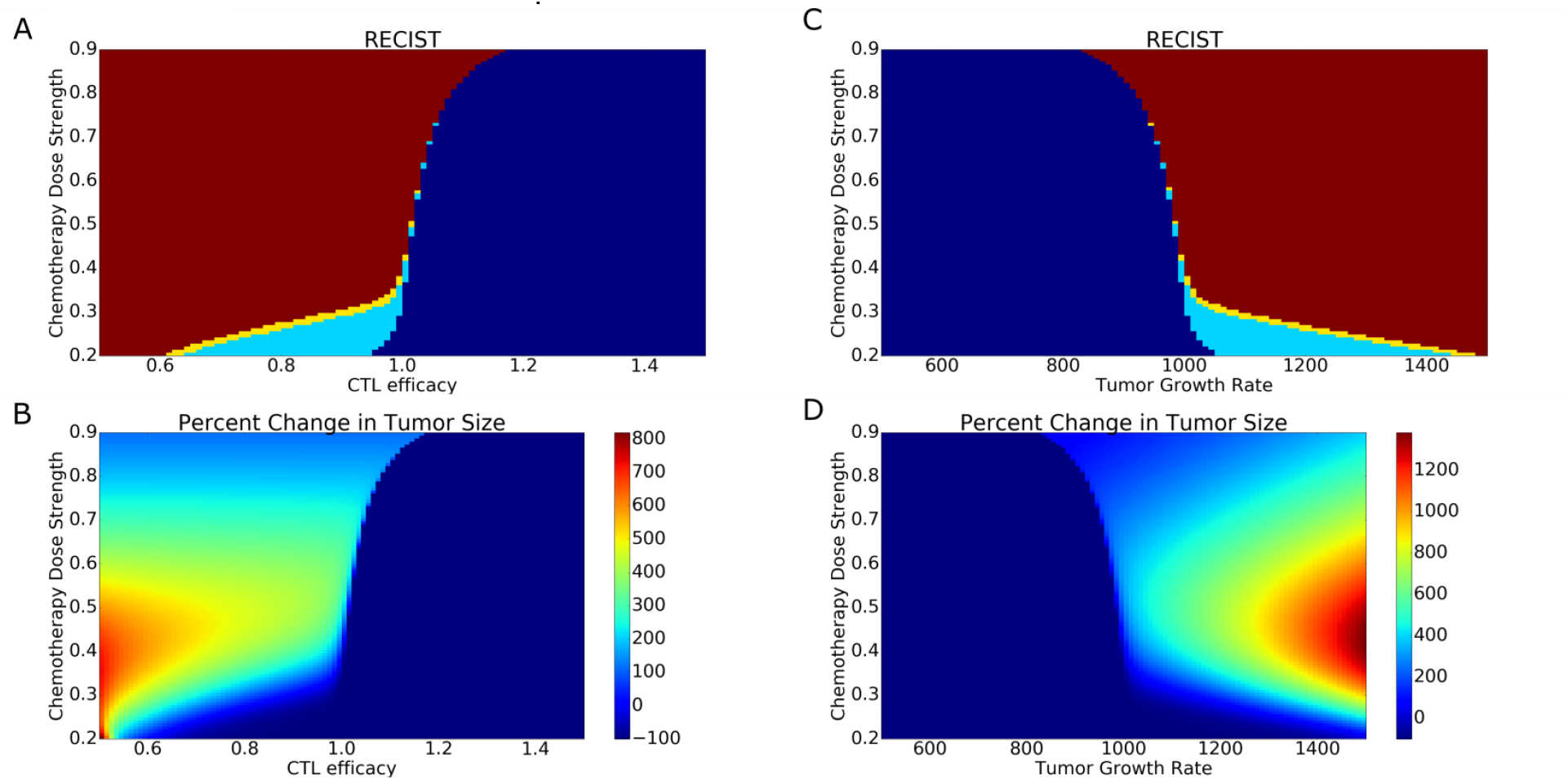
Treatment outcomes for variation in CTL efficacy (A and B) and tumor growth rate (C and D). Panels A and C represent RECIST outcomes. Red is progressive disease (PD), dark blue is complete response (CR), light blue is partial response (PR) and yellow is stable disease (SD). As CTLs become more efficient at killing tumor cells, there is a dramatic reduction in final tumor size and a significant improvement in outcome. However, below a threshold efficacy, chemotherapy has a much more important role in impacting the role of therapy. Weaker chemotherapy leads to better outcomes. A similar pattern is shown in response to variation in tumor growth rates. Faster growing tumors lead to significantly poorer treatment outcomes. This trend is most observable when, for chemotherapy values below 0.4, the range of tumor growth rates and CTL efficacies where tumor reduction is possible significantly increases. Chemotherapy plays an important modulating role in these faster growing tumors, however, with optimal treatment coming from weaker chemotherapy.

In short, patient immune biology determines optimal chemotherapy strength by determining which immune dynamics can be taken advantage of to control tumor growth. Low dose therapy is optimal in situations where the patient immune response is robust enough to control tumor growth. This requires both a sufficient memory-cell population as well as sufficiently high efficacy in CTL cytotoxicity. In contrast, high-dose chemotherapy is optimal to control tumor growth when either the immune system is unable to generate a sufficient CTL response, or when the tumor is slow-growing. However, in many situations where the immune system is able to enhance the effect of chemotherapy, dosing must be moderated so that it does not impose an overly large recovery burden and impede immune effects.

### Improvements to therapy outcomes from immunostimulatory vaccines: The Goldilocks Window

Patient-specific vaccines have become a recent hallmark in personalized cancer therapy. One of the first to acquire FDA approval was Sipuleucel-T, for treating metastatic castrate resistant prostate cancer [21]. Each vaccine is tailored to a specific patient by culturing dendritic cells from patient serum samples (taken roughly 72 hours before vaccine administration). The goal is to activate dendritic cells *in vitro* with a specific tumor protein target. These cultured antigen-presenting cells are then injected into the patient in order to stimulate an antitumor immune response. Three doses were administered in 2 week intervals with significant clinical responses being observed. Vaccination led to a 22% reduction in the relative risk of death, although there was no noticeable decrease in the rate of progression of disease [21]. The specific effect on T cells has been quantified by looking at T-cell receptor changes in response to vaccination. Subjects that received the vaccine saw a change in abundance and diversity of T-cell receptors in tumor-infiltrating lymphocytes. Certain receptor sequences were enriched, while others were significantly decreased [22], suggesting that the vaccine promoted an antigen-specific immune response against the tumor.

To study the effects and potential synergy of chemotherapy with this method of T-cell stimulation, we simulated a vaccine regime similar to that used for Sipuleucel-T (3 doses, spaced 14 days apart), with different vaccine strengths. Mathematically, this was modeled by modifying the ODEs that govern CTL expansion. The antigenicity parameter of the tumor, ***α***, was changed from a constant coefficient to a variable, time-dependent function, ***α_ν_***(***t***):

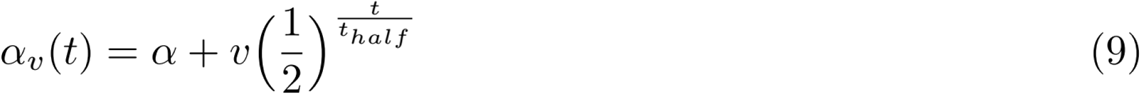

Total antigenicity is modeled as the result of both the constant, baseline antigenicity of the tumor, ***α***, and the exponentially decaying vaccine-augmented component, ***ν***. Vaccine-augmented antigenicity decays with a half-life, *t_half_*, of 3 days, a biologically realistic timespan in line with the short half-lives of dendritic cells [23]. This model of dynamic antigenicity can be expanded for multiple vaccinations, as used in the clinical protocol (eq. 10).

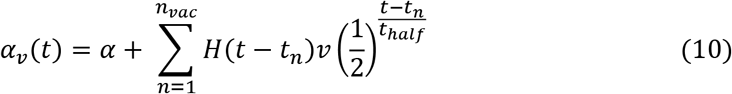

Here, H(t) is again the Heavyside function. *n_vac_* represents the total number of vaccine injections and *t_n_* represents the time of the n^th^ vaccination.

The ODEs used for the simulation of immune and tumor cell populations are then dependent on the instantaneous current value of ***α_ν_***(***t***) throughout the course of simulated therapy.

Under this scheme, results show that vaccine therapy can improve outcomes, but only within a specific range of chemotherapy strengths (Fig. 4). For very high chemotherapy doses, the beneficial effects of a vaccine are diminished. As before, the underlying cause for decreasing efficacy is the persistent lymphodepletion of memory cells due to the chemotherapy. Antigenicity augmentation due to vaccine stimulation is offset by reduced CTL expansion. However, very low-dose chemotherapy poses its own challenges, because with insufficient lymphodepletion, tolerogenic mechanisms and greater Treg recruitment inhibit any CTL response augmented by the vaccine. The immune system remains closer to tumor-tolerized homeostasis, and as a result vaccine stimulation is mitigated because the immune system is already suppressed.

**Figure 4:**
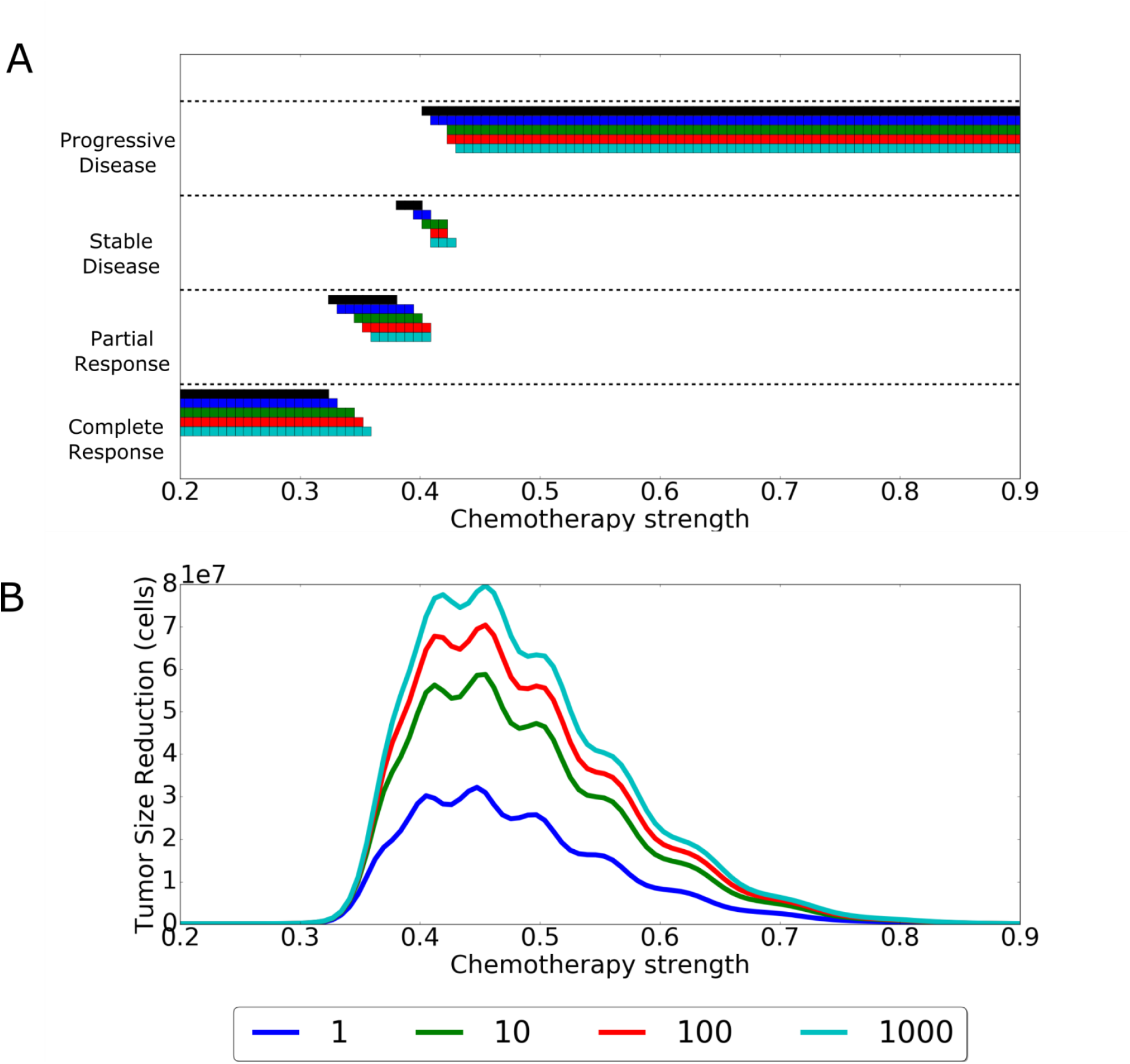
Improvements in tumor reduction due to vaccine application. Panel A shows the RECIST responses achieved for different vaccine strengths and chemotherapy strengths with black being the non-vaccine baseline. Vaccine strengths (***ν***) are 1 (blue), 10 (green), 100 (red), 1000 (light blue). Larger vaccine strengths lead to more successful RECIST responses for stronger chemotherapy doses. When looking at the absolute number of improvement in cellular reduction (B), a window of optimal chemotherapy ranges appears. Only when chemotherapy is in this range can vaccines provide a significant additional benefit.

Therefore, there exists an optimal dosing window for chemotherapy, a “Goldilocks” window. Quantitatively, we define this window to be the region in which a therapy dose can offer at least a 20% reduction in tumor size since this is the necessary amount for disease to become classified as a partial response. In order for there to be this maximized benefit from vaccine application, the chemotherapy regimen must be ‘just right’. Chemotherapy must have sufficient lymphodepletion to resensitize the immune system, but must leave enough immune cells such that vaccine stimulation leads to a large CTL response. Similar to the results of chemotherapy without the vaccine, the specific range of this Goldilocks window depends upon the initial patient memory cell (M_0_) numbers (not shown). More memory cells mean a system able to tolerate a larger dose of chemotherapy and still lead to a large vaccine-triggered CTL response. In contrast, fewer memory cells requires weaker chemotherapy doses to derive a maximum benefit from vaccine administration.

### Impact of variation in immune support

Chemotherapeutic lymphodepletion in the clinical setting can pose a serious threat to the safety of the patient through neutropenia [24], which commonly leads to dose reductions and disruptions to the standard schedule of therapy for patients. Consequently, multiple tools have been developed to help mitigate the effects of chemotherapy on the immune system. For example, it was recognized that dexamethasone treatment before carboplatin and gemcitabine could not only increase chemotherapy efficacy but also reduce the lymphodepleting effects by preventing uptake in the spleen and bone marrow [25]. In contrast, other aspects of cancer therapy can potentially hamper CTL responses to tumor insults. For example, G-CSF application has been shown to reduce CD8^+^ T cell activation and could conceivably impede the impact of lymphodepletion as a break from immune tolerance [26]. More generally, however, the broader impact of immune system augmentation or suppression during therapy remains unexamined.

In order to examine the effect of attenuated or augmented lymphodepletion on therapy outcome, we allowed for variable chemotherapeutic toxicity to immune populations, as compared to the tumor population. Mathematically, this simply means modifying the chemotherapy dose by a scaling factor **h**. The effect of chemotherapy on immune cell populations at a given treatment time is:

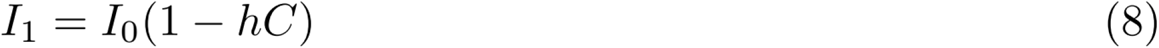

where ***I_1_*** is the immunological population size after application of chemotherapy, *I_0_* is the population size before therapy, and *0* < *C* < *1* is the dose strength. The specific numerical range in which *h* falls represents either attenuated or augmented chemotherapeutic toxicity. For values of *0* < *h* < *1*, this represents an attenuated toxicity relative to the toxicity on the tumor. In contrast, values of *h* > *1* represent higher toxicity on patient immune populations than on the tumor. This could be due to patient-dependent increased sensitivity to chemotherapy. However, this is really beyond the scope of our model, especially since mathematically I_1_ could become negative. This is clearly an area where our model may not accurately capture the dynamics. Therefore, we have restricted *hC* such that *hC* < *1*. For our *in silico* therapies, *h* was varied across these ranges where I_1_ > 0 for three different strengths of treatment. Values of *C* were chosen to represent lower (C= 0.25), middle (C= 0.6), and higher (C = 0.9) dose chemotherapy.

Outcomes of therapy due to variation in *h* depended upon the strength of chemotherapy. Interestingly, the results suggest that immune-supporting combination therapy has essentially no benefit when given with low dose chemotherapy. As shown in Figure 5, similar tumor reduction occurred for a wide range of values of *h* around *h*=*1* (which represents no immune support).

**Figure 5:**
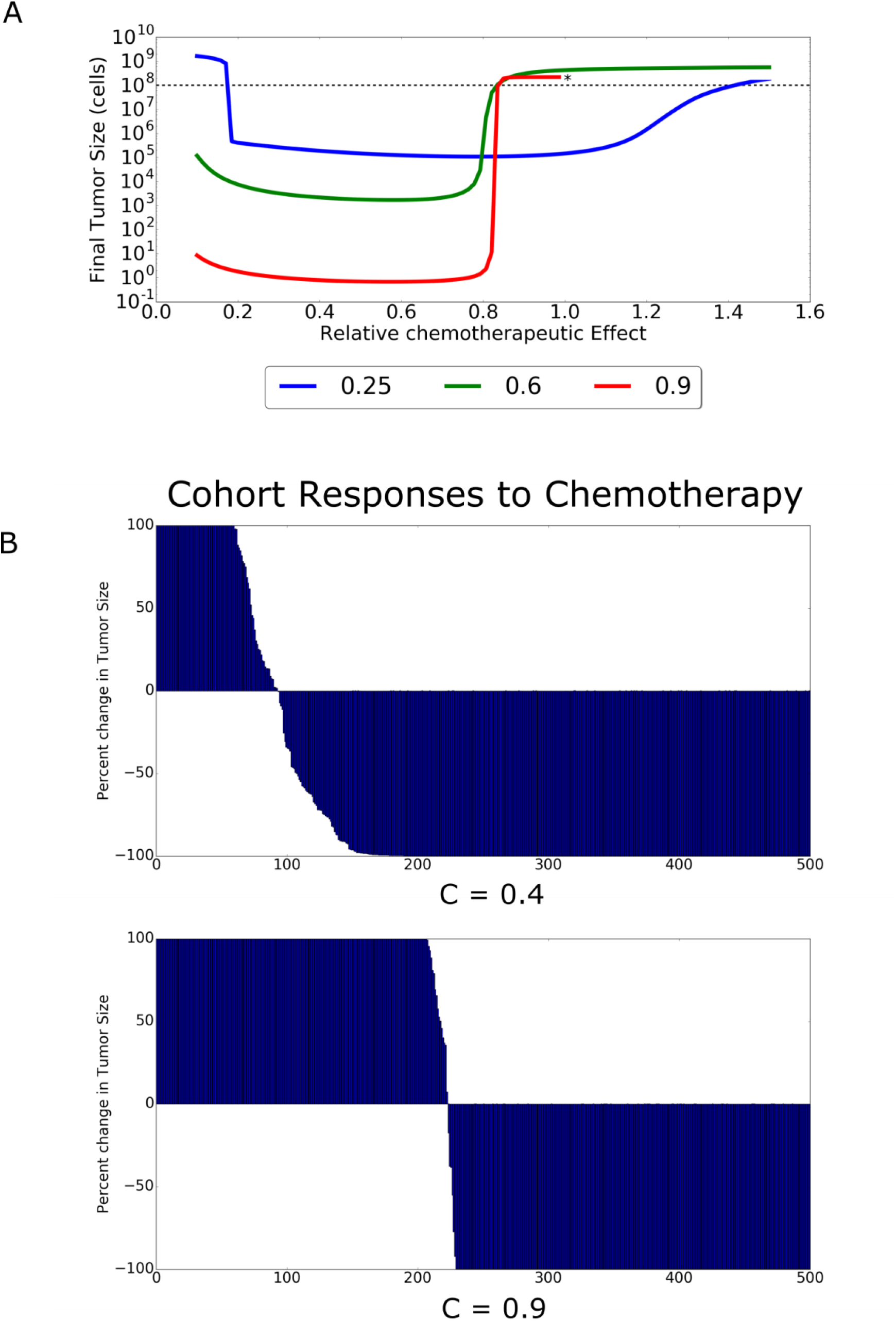
Therapeutic effects of differential response to immune prophylactics. (A) Final tumor sizes are shown for three different chemotherapy regimes for a range of immune modifier efficacies (**h**). The asterisk denotes that simulations were only run up to this ***h*** value for the highest dose chemotherapy. (B) Cohorts are treated with these differing regimes of high and low chemotherapy, showing significant differences in the proportion of successful versus unsuccessful responders.

Furthermore, outcomes were worse when *h* was very low or very high. In situations where it was very low, final tumor sizes were large because a lack of lymphodepletion did not sufficiently break immune tolerance. In contrast, for larger *h* values, there was over-depletion which prevented an effective T-cell response despite significant tolerance breaking.

In contrast, high dose chemotherapy saw treatment failure or success highly dependent upon the amount of immune support. Similar to low dose therapy, a small value of *h* that mitigated the depleting effects of chemotherapy led to the best possible outcomes in terms of tumor shrinkage. Final tumor sizes were, in fact, multiple orders of magnitude lower than was possible with low-dose chemotherapy. As ***h*** increased (representing less toxicity mitigation) treatment outcomes rapidly worsened. The transition value ***h\****, where the clinical outcome rapidly shifts, indicates a threshold effect with regard to immune support. For high chemotherapy doses, immune support treatments must have a significantly large mitigation (***h*** < ***h\****) of immunodepletion in order for successful treatment responses to occur.

Interestingly, the moderate strength chemotherapy regimen yielded only partial benefits of either extreme. The greatest tumor reduction possible, with immune support, yielded tumors that were smaller than those achievable with low dose chemotherapy. However, these tumors were still multiple orders of magnitude larger than those achievable with high dose chemotherapy. For treatment failure at lower immune support (***h*** > ***h\****) tumor sizes were actually larger than when high dose chemotherapy failed.

Clinically, the results suggest that chemotherapy dose strength can be used to mitigate uncertainty regarding the amount of immune support a certain treatment will give to a specific patient. Low dose therapy offers a wide range of potential immune support in which treatment can successfully reduce tumor sizes. The disadvantage is that the maximum tumor size reduction still leaves larger tumors than are possible using higher doses of chemotherapy. While our model has not analyzed this, a potential impact is that larger tumor sizes could lead to more heterogeneous populations and thus lead to a higher likelihood of resistant or metastatic populations. However, higher doses have a narrower range of immune support in which they are successful. Chemotherapy can be balanced, then, against how certain the clinician is of the benefit that G-CSF (or other immune supporting drug) will give. For patients where there is high certainty of a significant benefit due to the drug, high dose therapy is optimal. In contrast, lower dosing should be used when the drug may have lower or variable efficacy.

### Variable Immune Support and Impacts on Observed Cohort Responses

Finally, we sought to investigate how variation in the effectiveness of these immune adjuvants might impact treatment outcomes in a group of patients. Chemotherapy treatment leads to a wide range of responses, both successful and unsuccessful, across multiple types of cancer [17]. This variation has been attributed to both disease variation, patient variation, and interactions between the two. However, less attention has been given to variable patient responses to secondary drugs – such as G-CSF – and how they impact therapy. Patient responses to these secondary drugs are currently poorly measured and could have significant implications for therapy outcomes.

To better explore the effect of variable patient responses to immune support drugs, cohorts of 500 patients were randomly generated from a normal distribution with a mean immune support response value of ***h*** = 0.8 and variance of 0.2. These values were chosen to center the distribution around the model-derived threshold value ***h* = 0.8***. Similar to our previous investigations, cohorts were then subjected to regimens of low (C = 0.4) and high (C = 0.8) chemotherapy strengths. Percent changes in tumor size after therapy were displayed for each individual patient in the cohort to generate a waterfall plot. In doing so, we used our model to simulate cohort responses as is commonly measured in aggregated studies of patient data [17]. The waterfall plots (Fig. 5) illustrate that chemotherapy strength can significantly change the proportion of successfully responding patients in a population with variable responses to immune prophylactics. This is significant since the proportion of successful responses is often an important criterion for judging therapeutic efficacy. The simulated waterfall plots show how clinical outcomes could not only be the result of therapy, but also due to inherent immune variation within the cohort.

## Discussion

A major barrier to success for immunotherapy in cancer is tolerogenic mechanisms that reduce the immune response to tumor antigens ([27], [3], [6]). A potential solution has come from observations that lymphodepletion stimulates homeostatic proliferation in the immune system which can transiently restore immune response. This has led to increasing efforts to selectively apply chemotherapy to improve outcomes from immunotherapy [28].

To better understand this potential synergy, we constructed a mathematical model to frame these complex dynamics and identify critical parameters that govern the clinical outcomes. Our studies focused on three clinically-observed dynamics of immunodepletion, immunostimulatory vaccination, and immunosupportive prophylactics. With regard to immunodepletion, we demonstrated that chemotherapy results in a trade-off. At very high doses, chemotherapy has a maximal cytotoxic effect on the tumor but also maximally depletes memory T cells such that no effective CTL response can be mounted despite the transient loss of tolerance during re-expansion of the immune cells after completion of chemotherapy. Similarly, low doses of chemotherapy are insufficient to produce the post-treatment immune cell expansion that is necessary for reversal of immune tolerance. Importantly, however, we find there is a “Goldilocks” range of chemotherapy doses in which lymphodepletion causes adequate immune resensitization, but does not impose an overly large recovery burden. This window is governed by the patient-specific quantity of memory T cells so that larger pre-treatment T-cell populations allow more favorable outcomes with higher doses of chemotherapy. In contrast, fewer pretreatment CTLs can limit the immune response even in the “Goldilocks” range of chemotherapy. Thus, there is a necessary ‘minimum efficacy’ of effector cells for successful stimulation of immune response by chemotherapy. Below this threshold of immune activity, the benefit of chemotherapy is almost solely dependent on its inherent cytotoxicity (Fig. 6)

**Figure 6:**
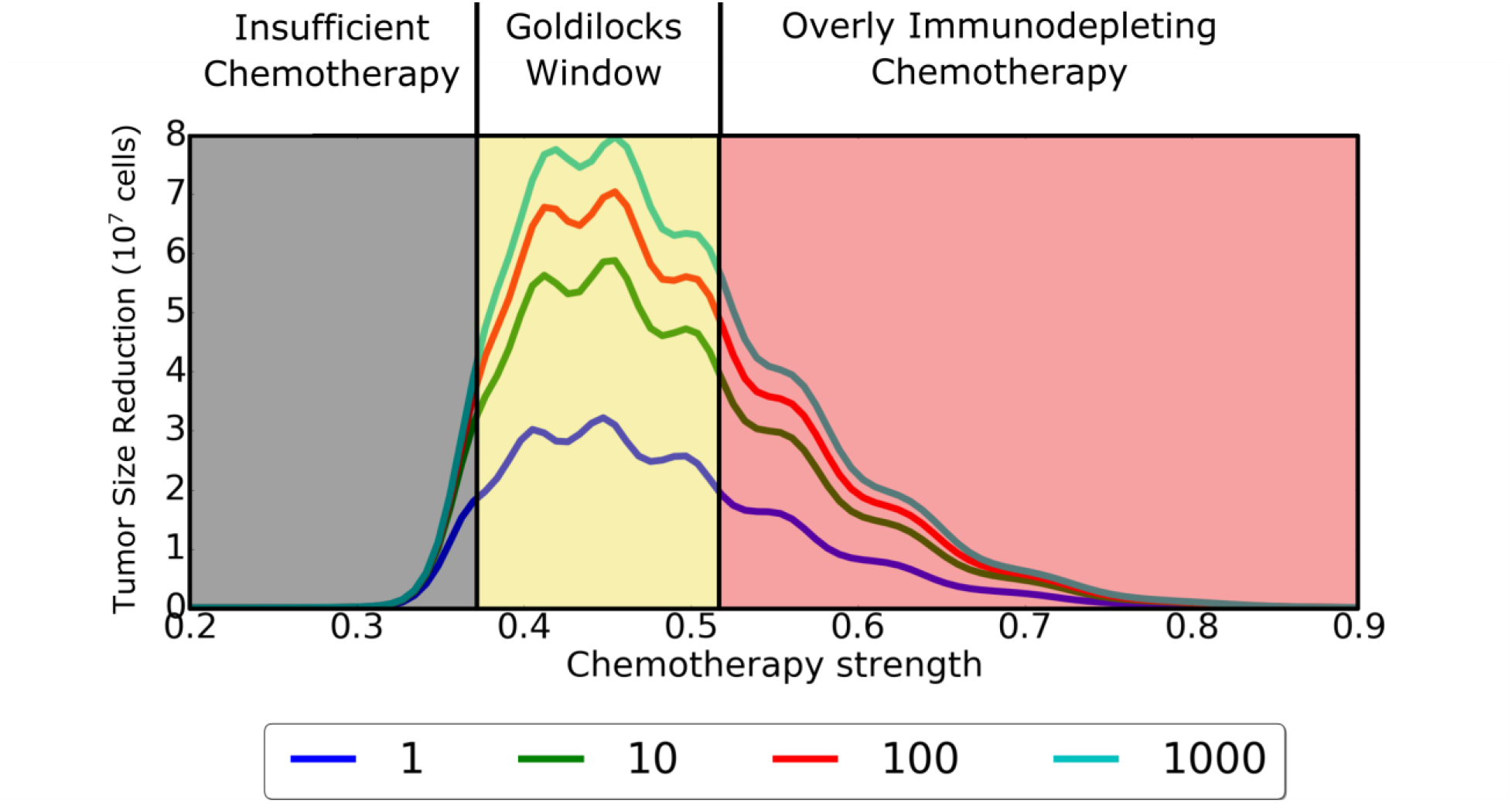
A diagram explaining tumor outcomes at varying chemotherapy strengths and immune support doses. If therapy is too weak, then immune stimulation cannot be maximally effective and direct chemotherapy-mediated tumor cell death is also low. This yields a suboptimal tumor reduction. When chemotherapy is too strong, there may be more tumor cell death due to the drug, but insufficient immune activation due to over depletion of T cells. There is a moderate dose, however, that represents a Goldilocks window of maximizing both T-cell activation as well as drug-induced tumor cell death. This range of dosing provides at least a 20% reduction in tumor size (relative to the initial tumor size of 10^8^ cells).

Our model also provides insight into the potential effects of variation in the tumor growth rate. In slower growing tumors, chemotherapy alone can be sufficient to achieve optimal treatment response. Treatment of faster growing tumors, however, is best when the chemotherapy is administered to enhance the immune response. Unfortunately, if the pre-treatment population of CTLs is small, we find chemotherapy for rapidly growing tumors will be ineffective if it is both highly lymphodepleting and insufficiently cytotoxic to significantly reduce tumor growth. Assessing the clinical importance of this question is challenging because it remains unclear from the literature as to the actual size of the population of tumor-specific T cells that are present during treatment. In spite of these difficulties, the impact and existence of anti-tumor immunity has been bolstered by recent immunotherapies which act to remove inhibitions to T-cell action [29].

Chemotherapy is increasingly being used in concert with vaccines to help stimulate the patient immune system. We investigated the interactions between vaccines and lymphodepletion and found that, as before, there is a window of chemotherapy ranges in which vaccines can improve outcomes versus chemotherapy alone. At very high doses, however, the resulting lymphodepletion substantially reduces benefits of immune stimulation by vaccination. More broadly, other novel immunotherapies could also potentially be hampered by over-depletion of the immune system.

To further investigate the potential impact of this interaction, we modeled the effect of differential responses to immune prophylactics. G-CSF and other drugs have become common recourses in chemotherapy for mitigating the immunodepletion effects on patients [30]. However, recent studies have suggested that T cell response is hampered by G-CSF administration [26]. While G-CSF may help prevent neutropenia and cytopenia for patients, it may impede the ability of retolerized T cells to mount an anti-tumor response. In addition, responses to prophylactics are not constant but the significance of this variation remains relatively uninvestigated. Our model suggests that inter-patient variation in prophylactic response can lead to drastically different outcomes for the same dosing of chemotherapy. Across larger samples, this variation can further interact with chemotherapy to be a significant determinant of whether the chemotherapy dose leads to more success or failure across a range of patients.

In conclusion, our results suggest opportunities to increase the efficacy of immunotherapy with precise application of chemotherapy. Our model affirms the importance of effector and memory T-cell expansion following chemotherapy to reduce immune tolerance to tumor antigens. However, we demonstrate that optimal chemotherapy requires identification of a Goldilocks Window in which treatment can both induce cytotoxic effects in the tumor and enhance the immune response to tumor antigens. Identifying optimal strategies for chemotherapy in each patient will likely benefit from the application of mathematical models which are parameterized by patient data pre-treatment to generate an optimal treatment strategy for that patient. Importantly, these predicted strategies would most likely need to change as patient responses diverge from those predicted, leading to an iterative loop of ‘predict-apply-refine’. With the growing drive towards precision medicine, we believe that mathematical models are critical for the future of truly personalized therapy, where no two patients will receive the same therapeutic regimen, and where treatments adapt a change based on patient responses. The model presented here is a step towards describing the complex landscape of treatment decisions regarding dosing and combination of different therapies, and we have shown how these decisions can be sensitive to patient-specific parameters and guide clinical intuition.

## Notes

**Conflict of Interest statement**: The authors declare no potential conflicts of interest.

